# *tarsal-less* is expressed as a gap gene but has no gap gene phenotype in the moth midge *Clogmia albipunctata*

**DOI:** 10.1101/266288

**Authors:** Eva Jiménez-Guri, Karl R. Wotton, Johannes Jaeger

**Author notes:** Current address: Centre for Ecology and Conservation, College of Life and Environmental Sciences, University of Exeter, Penryn, Cornwall TR10 9EZ, UK. Current address: Complexity Science Hub Vienna, Josefstädter Straße 39, 1080 Vienna, Austria, and Center for Systems Biology Dresden (CSBD), Pfotenhauerstrasse 108, 01307 Dresden.

## Abstract

Gap genes are involved in segment determination during early development of the vinegar fly *Drosophila melanogaster* and other dipteran insects (flies, midges, and mosquitoes). They are expressed in overlapping domains along the antero-posterior (A–P) axis of the blastoderm embryo. While gap domains cover the entire length of the A–P axis in *Drosophila*, there is a region in the blastoderm of the moth midge *Clogmia albipunctata*, which lacks canonical gap gene expression. Is a non-canonical gap gene functioning in this area? Here, we characterize *tarsal-less* (*tal*) in *C. albipunctata*. The homolog of *tal* in the flour beetle *Tribolium castaneum* (called *milles-pattes, mlpt*) is a *bona fide* gap gene. We find that *Ca-tal* is expressed in the region previously reported as lacking gap gene expression. Using RNA interference, we study the interaction of *Ca-tal* with gap genes. We show that *Ca-tal* is regulated by gap genes, but only has a very subtle effect on *tailless (Catll)*, while not affecting other gap genes at all. Moreover, cuticle phenotypes of *Ca-tal* depleted embryos do not show any gap phenotype. We conclude that *Ca-tal* is expressed and regulated like a gap gene, but does not function as a gap gene in *C. albipunctata*.

## Introduction

The gap gene network provides the first layer of zygotic regulation in the segmentation gene hierarchy of dipteran insects (flies, midges, and mosquitos). In the vinegar fly *Drosophila melanogaster*, this network consists of the trunk gap genes *hunchback (hb), Krüppel (Kr), knirps (kni)*, and *giant (gt)*, with additional inputs from the terminal gap genes *tailless (tll)* and *huckebein (hkb)* [1]. In other cyclorrhaphan flies, such as the hoverfly *Episyrphus balteatus* [2,3] and the scuttle fly *Megaselia abdita* [4–6], gap gene expression and regulation is strongly conserved. Outside the cyclorrhaphan clade, among the nematoceran Diptera, there is little functional evidence on gap gene regulation although expression patterns have been described in the malaria mosquito *Anopheles gambiae* [7].

Here, we focus on another emerging nematoceran model system, the moth midge *Clogmia albipunctata* (Diptera, Psychodidae). In this species, we have a detailed description of the spatial arrangement [8,9] as well as the temporal dynamics [10] of gap gene expression. This descriptive evidence reveals a region of the *C. albipunctata* blastoderm embryo, which is not covered by expression of any gap gene known from *D. melanogaster* [9]. This region lies between the abdominal domain of the *C. albipunctata* homolog of *kni* and *knirps-related* (called *knirps-like, knl*) and the posterior terminal domain of *tll* [9,10]. It suggests that we may be missing a posterior gap gene in this species.

One candidate for this missing gap gene in *C. albipunctata* is *tarsal-less (tal)* [11], also called *polished rice (pri)* [12]. *tal/pri* is a polycistronic gene encoding a long primary transcript from which several short peptides are produced that are required in different stages of embryonic development. It is part of a large class of polycistronic genes with small open reading frames (sORF/smORF), small encoded peptides (SEP), or microproteins that play a wide range of roles in physiology, development, and cell differentiation [13,14]. In *D. melanogaster, tal/pri* is first expressed in a stripe-like expression pattern at the late blastoderm stage [12]. It is involved in epithelial morphogenesis and leg development [11,12,15–18], but has no role in early embryonic patterning or segment determination.

Interestingly, a homolog of *tal/pri* was first described in the flour beetle *Tribolium castaneum* under the name of *mille-pattes* (*mlpt*) [19]. In contrast to *tal/pri* in *D. melanogaster, mlpt* in *T. castaneum* has a segmentation function acting as a *bona fide* gap gene [19]. *mlpt* is expressed in a gap-like fashion, with an anterior and a posterior terminal domain at blastoderm stage; subsequently, the anterior domain resolves into two stripes, and the terminal domain retracts from the pole and shifts anteriorly over time during germband extension; a third posterior domain appears at this stage; finally, *mlpt* is expressed in the peripheral nervous system and the forming appendage joints at later stages of development, which is similar to its expression pattern in *D. melanogaster* [19]. Knock-down of *mlpt* in *T. castaneum* by RNA interference (RNAi) leads to a gap-like phenotype with missing abdominal segments [19]. *mlpt* regulates trunk gap genes *hb, Kr*, and *gt*, and is itself regulated by *hb* and *Kr* [19].

Here, we characterize expression of *tal/pri* in *C. albipunctata*, and examine its interactions with other segmentation genes using RNAi knock-down assays. We show that it exhibits a gap-gene-like expression pattern at the blastoderm stage. As in *T. castaneum*, it is expressed in an anterior and a posterior terminal domain, which later split into narrow stripes. In contrast to *T. castaneum*, however, *tal/pri* does not regulate gap genes in *C. albipunctata*, with the possible exception of its interaction with the posterior terminal *tll* domain. Even though it is regulated by gap genes *hb, Kr*, and *knl*, it does not exhibit any gap-like phenotype when knocked down. This evidence suggests that although *tal/pri* is expressed and regulated in a gap-gene-like manner, it cannot be classified as a *bona fide* gap gene in *C. albipunctata*.

## Results and Discussion

### *Characterisation of* tarsal-less *in the moth midge* C. albipunctata

We searched the early embryonic transcriptome of *C. albipunctata* [20] for a *tal* homologue using the *D. melanogaster* amino acid sequences for the small peptides encoded by *tal*. Our search identified a 2277 nt fragment that contained several short peptide repeats, probably corresponding to a primary transcript. Upon *in silico* translation, it was confirmed as a homolog of *tal/pri/mlpt* in *C. albipunctata*. We will call this fragment *Ca-tal*. Specific primers were generated to clone the gene from cDNA, and empirically confirm its sequence (see Materials and Methods). The *Ca-tal* sequence has been deposited in GenBank under accession number MG783326.

The polycistronic sequence of *Ca-tal* shows general structural similarities to *tal* genes in other organisms (Figure 1A). *tal* genes exhibit variable numbers of repeats of N-terminal peptides containing a consensus region of LDPTGXY, and one C-terminal peptide with the consensus domain GREETSSCRRRR [19]. In *Ca-tal*, we find four short repeated N-terminal peptides of 11, 10, 11, and 29 amino acids separated by 111, 126, and 334 nt length respectively, plus a longer 39 amino acid C-terminal peptide separated from the closest N-terminal peptide by 348 nt (Figure 1B).

**Figure 1.**
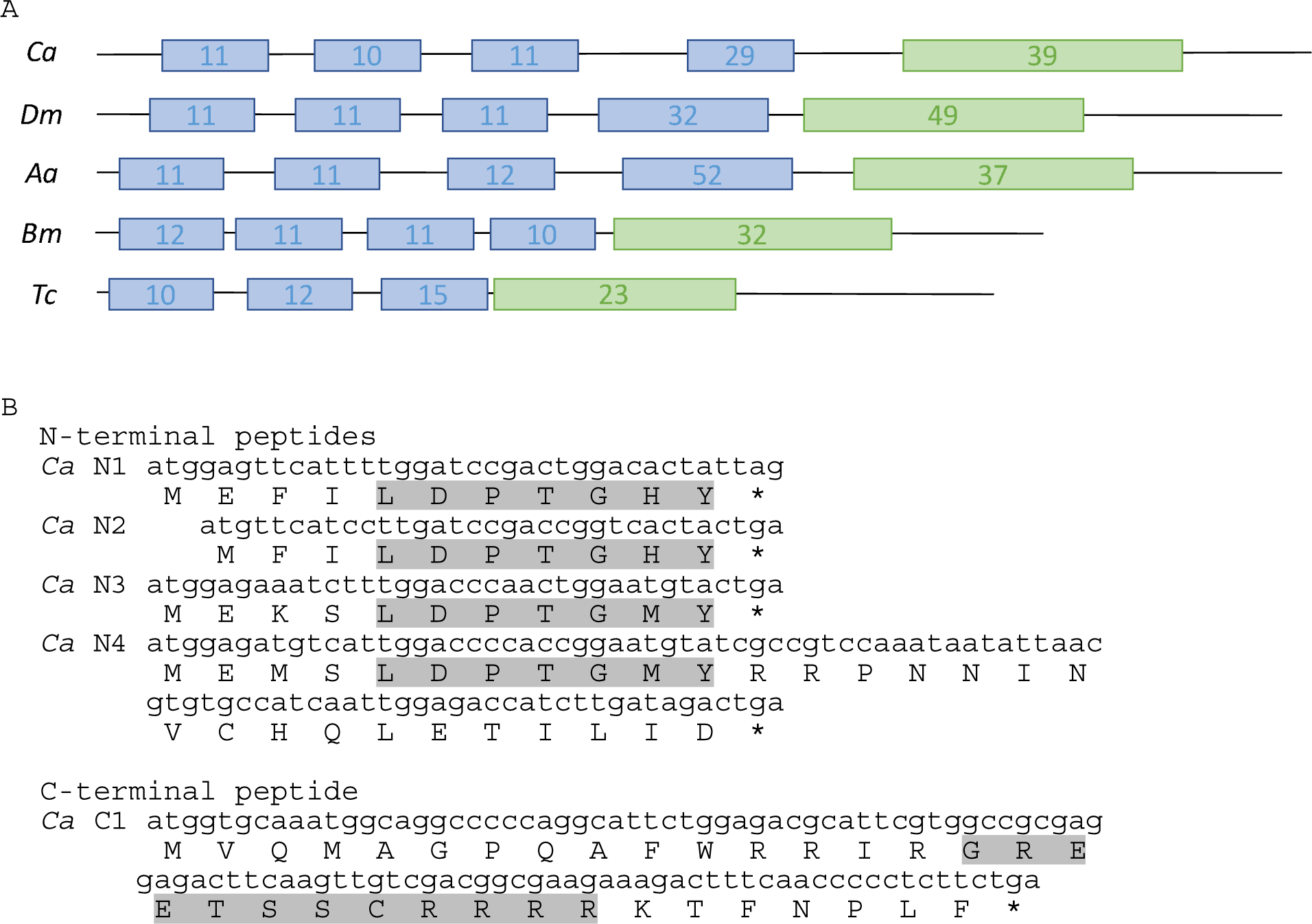
Sequence of *Ca-tal* and *in silico* translation. (A) Schematic comparisons of small ORFs in *tal/pri/mlpt* homologs of *C. albipunctata (Ca), D. melanogaster (Dm), Aedes aegypti* (*Ae*), *Bombyx mori* (*Bm*) and *T. castaneum (Tc)* (sequences as described in [19]). Blue boxes represent N-terminal peptides; green boxes, C-terminal peptides. Numbers represent peptide length in amino acids. The fourth N-terminal peptide in *D. melanogaster* and *A. aegypti* have two conserved repeats. (B) Five conserved peptides are found in the transcriptomic sequence of *Ca-tal*, as retrieved from the Diptex database (CAL_comp2583_c0_seq1): four N-terminal (*Ca* N1-4), and one C-terminal peptide (*Ca* C1). Conserved consensus regions for each peptide types are highlighted in grey. Complete primary transcript sequence is deposited in GenBank with accession number MG783326.

### *Temporal expression profile of* Ca-tal *in the embryo*

We have characterised the expression pattern of *Ca-tal* in the embryo of *C. albipunctata* from the blastoderm up to the extended germband stage [21] using enzymatic (colorimetric) *in situ* hybridization (Figure 2). The earliest pattern we detect is a posterior expression domain in the trunk region of the blastoderm embryo, covering 65–80% anteroposterior (A–P) position (Figure 2A). This domain shifts anteriorly over time (Figure 2B). By the time it has reached 55–75% A–P position, a second terminal domain becomes apparent at the posterior pole (Figure 2C). Both domains continue to shift and expand anteriorly (Figure 2D), consistent with shifts observed for posterior gap genes during the blastoderm stage [9,10]. Before gastrulation, the anterior border of the more anterior *Ca-tal* domain reaches 55% A–P position (Figure 2D, arrowhead), and this domain starts to split into two stripes (Figure 2D, asterisks). By the same time, the posterior terminal domain has expanded to 85% A–P position.

**Figure 2.**
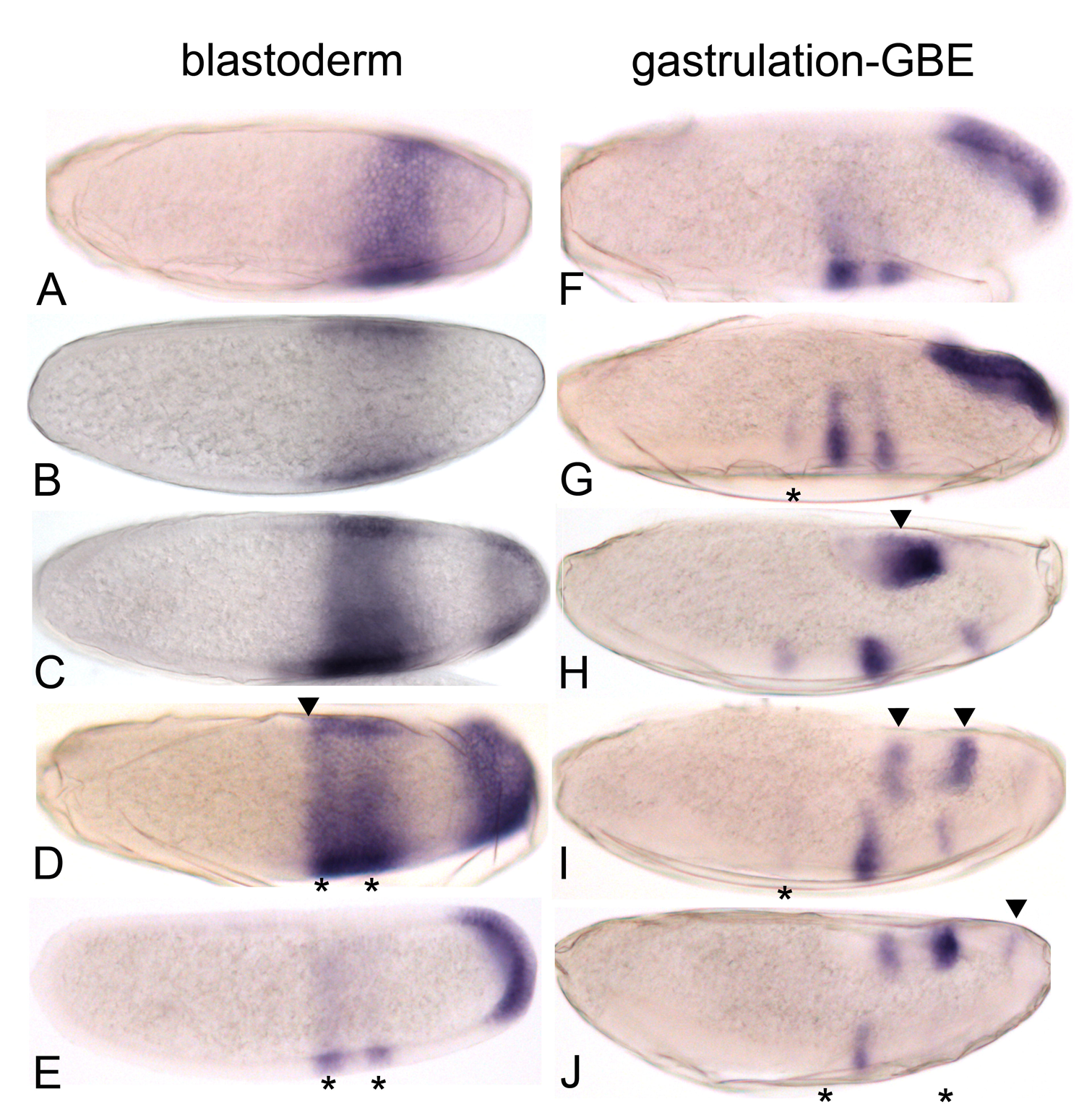
Expression pattern of *Ca-tal* during early embryo development of *C. albipunctata*. Colorimetric (enzymatic) *in situ* hybridisation against *Ca-tal* (blue) in embryos ranging from early cellular blastoderm (A) to gastrulation (F), and germ band elongation (GBE) (G–L). Time increases downwards in each column. Embryos oriented laterally: anterior is to the left, dorsal to the top. See text for details.

By the onset of gastrulation, the anterior domain has resolved completely into two stripes (Figure 2E, asterisks). A weak third stripe appears shortly thereafter in a more anterior position (Figure 2G, asterisk). This dynamic pattern is similar to what has been reported for the *tal* homolog *mille-pattes (mlpt)* in the flour beetle *T. castaneum* [19]. The terminal domain follows the morphogenetic movement of the posterior pole region during gastrulation [21], moving to the dorsal side of the embryo (Figure 2F–H); at the same time, this domain clears from the pole (Figure 2H, arrowhead) and divides into two sub-terminal stripes (Figure 2I, arrowheads). During germband elongation, the first, and later the third, stripe of the anterior domain fade away (Figure 2I,J, asterisks). Finally, an additional stripe appears anterior of the two sub-terminal stripes (Figure 2L, arrowhead).

As a next step, we performed double *in situ* hybridizations to define the expression of *Ca-tal* in reference to the gap gene domains in the *C. albipunctata* blastoderm (Figure 3). The anterior border of the more anterior *Ca-tal* domain coincides with the posterior border of the anterior domain of *Ca-hb*, both shifting anteriorly in concert over time (Figure 3A). The central domain of *Ca-Kr* and the more anterior *Ca-tal* domain show extensive overlap, although the latter extends slightly further posterior (Figure 3B). Domains of *Ca-gt* and *Catal* never overlap and are positioned far from each other in the embryo (Figure 3C). This is because *C. albipunctata* lacks a posterior *gt* domain, unlike *D. melanogaster* [9]. The more anterior domain of *Ca-tal* and the abdominal domain of *Ca-knl* overlap in the anterior region of the latter (Figure 3D). The posterior border of the abdominal *Ca-knl* domain coincides with the anterior border of the terminal *Ca-tal* domain (Figure 3D). The posterior terminal *Ca-tll* domain overlaps with the posterior part of the terminal *Ca-tal* domain (Figure 3E). In our double *in situs*, we see that the terminal *Ca-tal* domain already clears from the posterior pole during the blastoderm stage, so that its posterior boundary comes to coincide with the anterior border of *Ca-tll* just before the onset of gastrulation (Figure 3E).

**Figure 3.**
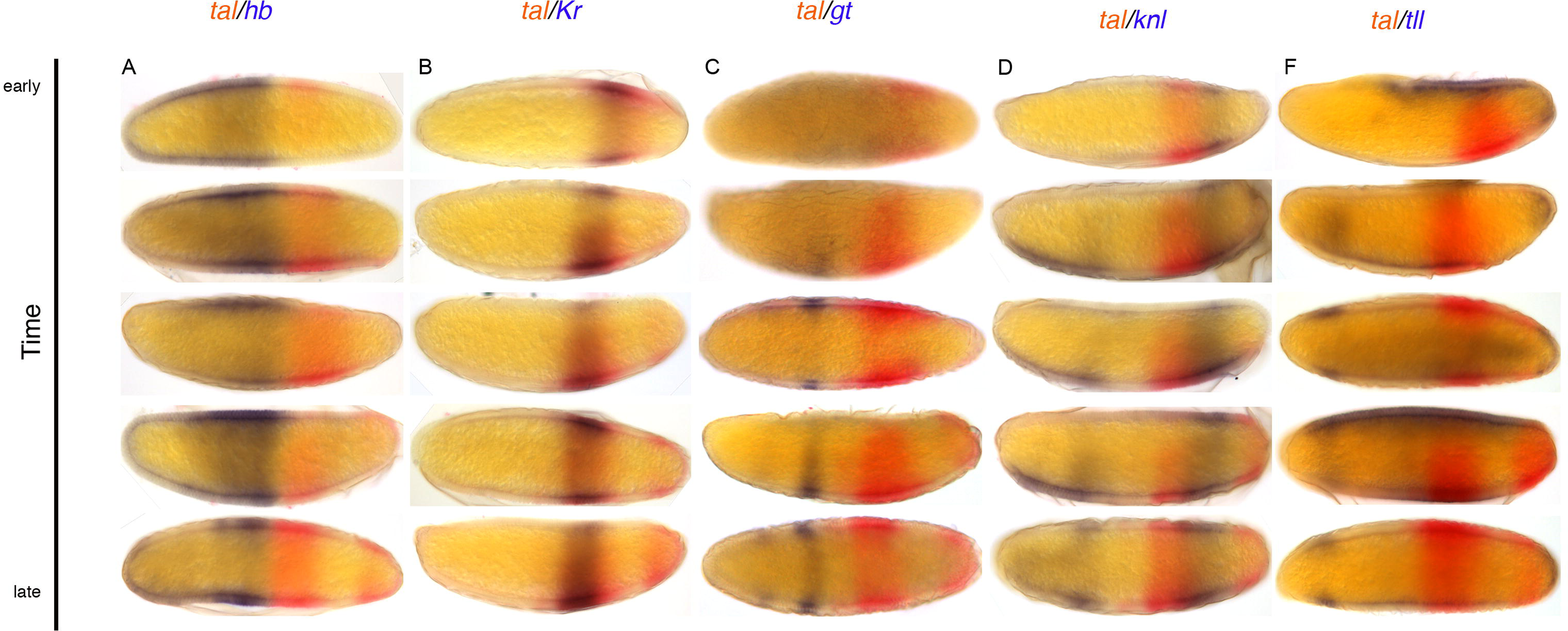
Relative localisation of *Ca-tal* and gap gene expression in *C. albipunctata* blastoderm embryos. Colorimetric (enzymatic) *in situ* hybridisation against *Ca-tal* (red) is shown with RNA patterns of gap genes (in blue) during the blastoderm stage. Stains as indicated by column headings: (A) *tal* (red)/*hb* (blue); (B) *tal* (red)/*Kr* (blue); (C) *tal* (red)/*gt* (blue); (D) *tal* (red)/*knl* (blue); (E) *tal* (red)/*tll* (blue). Time increases downwards. Embryos oriented laterally: anterior is to the left, dorsal to the top.

Our results show that *Ca-tal* is expressed in a gap-gene-like manner during the blastoderm stage, partially overlapping with previously characterized gap domains in *C. albipunctata* [9]. Intriguingly, the terminal *Ca-tal* domain covers a region of the *C. albipunctata* blastoderm—between the abdominal *Ca-knl* domain and the terminal domain of *Ca-tll*—in which no gap gene expression has been detected before [9,10]. In contrast, *tal* is not expressed like a gap gene in *D. melanogaster*, where its transcripts appear directly in a stripe-like pattern during the late blastoderm stage [12]. Early *Ca-tal* expression shows much more resemblance to that of its homolog *mlpt* in *T. castaneum*, which acts as a *bona fide* gap gene in that species [19]. This suggests that *Ca-tal* may also play the role of a gap gene in *C. albipunctata.* In order to test this possibility, we performed knock-down by RNA interference (RNAi) of *Catal, Ca-tll*, and other trunk gap genes.

### Ca-tal *does not regulate, but is regulated by trunk gap genes*

To assess the effect of *Ca-tal* on the gap genes in *C. albipunctata*, we performed RNAi knock-down against *Ca-tal* following a previously published protocol [22]. The resulting *tal*-depleted embryos were stained by colorimetric *in situ* hybridisation (ISH) for trunk gap genes *Ca-hb*, *Ca-Kr*, Ca-*gt*, and *Ca-knl*, as well as the terminal gap gene *Ca-tll.* The other terminal gap gene, *huckebein* (*hkb*), is not expressed at the relevant stages in *C. albipunctata* [9]. We do not observe any clearly detectable differences in the expression patterns of the trunk gap genes in *Ca-tal* knock-down embryos (Supplementary Figure 1). Quantitative assessment of domain boundary positions using our FlyGUI/FlyAGE image-processing pipeline [10,23] does not reveal any significant differences to the wild-type either (not shown). The only potential effect of *Ca-tal* on gap genes is the reduced expression in the posterior terminal domain of *Ca-tll* in a small percentage of *Ca-tal* knockdown embryos (4 out of 17; Supplementary Figure 1E,F; K,L). Target genes further downstream in the segmentation gene cascade, such as the pair-rule gene *even-skipped (eve)*, and the segment polarity genes *wingless (wg)* and *engrailed (en)*, also fail to show any clearly detectable defects in *Ca-tal* knock-down embryos (not shown). This suggests that *Catal* does not play any essential role in segmentation gene regulation in *C. albipunctata.*

Next, we investigated whether *Ca-tal* is regulated by gap genes. We assayed *Ca-tal* expression in embryos treated with RNAi against *Ca-hb*, *Ca-Kr*, *Ca-gt*, *Ca-knl*, and *Ca-tll* using colorimetric *in situ* hybridization (Figure 4). The expression pattern of *Ca-tal* was affected by all gap genes with the exception of *Ca-gt* (not shown). In blastoderm embryos depleted of *Ca-hb*, the more anterior domain of *Ca-tal* is displaced anteriorly, extending past 45% A–P position (Figure 4A; 10 out of 28 embryos). This suggests that Ca-Hb positions the anterior border of expression of *Ca-tal* through repression. In embryos depleted of *Ca-Kr*, we observe a loss of the more anterior *Ca-tal* domain, while its terminal domain appears to expand anteriorly (Figure 4B, 17 out of 20 embryos). This is consistent with a dual influence of Ca-Kr, with an activating effect on the more anterior domain, and repression on the terminal domain of *Ca-tal*. However, it is not clear whether both of these effects are direct. Activation could be mediated through repression of repressor *Ca-knl* by Ca-Kr. This is unlikely, as *Ca-knl* is not affected in knock-downs of *Ca-Kr* (Supplementary Figure 2, 17 out of 17 embryos). Still, we cannot exclude indirect activation mediated through repression of another unknown repressor. Finally, the effect of *Ca-Kr* could be interpreted as a deletion of the region between the two *Ca-tal* domains. This, however, seems unlikely, since *Ca-Kr* is not expressed near the potentially affected region of the embryo (cf. Figure 3B) and *Ca-knl* is still expressed there in *Ca-Kr* RNAi-treated embryos (Supplementary Figure 2B). In embryos depleted of *Ca-knl*, we see strong ectopic expression of *Ca-tal* between its two domains of expression at the blastoderm stage (Figure 4C, 25 out of 54 embryos). This suggests repression *of Ca-tal* by Ca-Knl. The effect is probably weak, since the de-repression seen in Figure 4C is incomplete, and the expression patterns of *Ca-tal* and *Ca-knl* show extensive overlap in the wild-type (Figure 3D). In late blastoderm embryos depleted of *Ca-tll*, the terminal domain of *Ca-tal* expression is either completely abolished or strongly reduced (Figure 4D, 17 out of 44 embryos). Taken together, our evidence suggests that *Catal* and *Ca-tll* activate each other in *C. albipunctata*.

**Figure 4.**
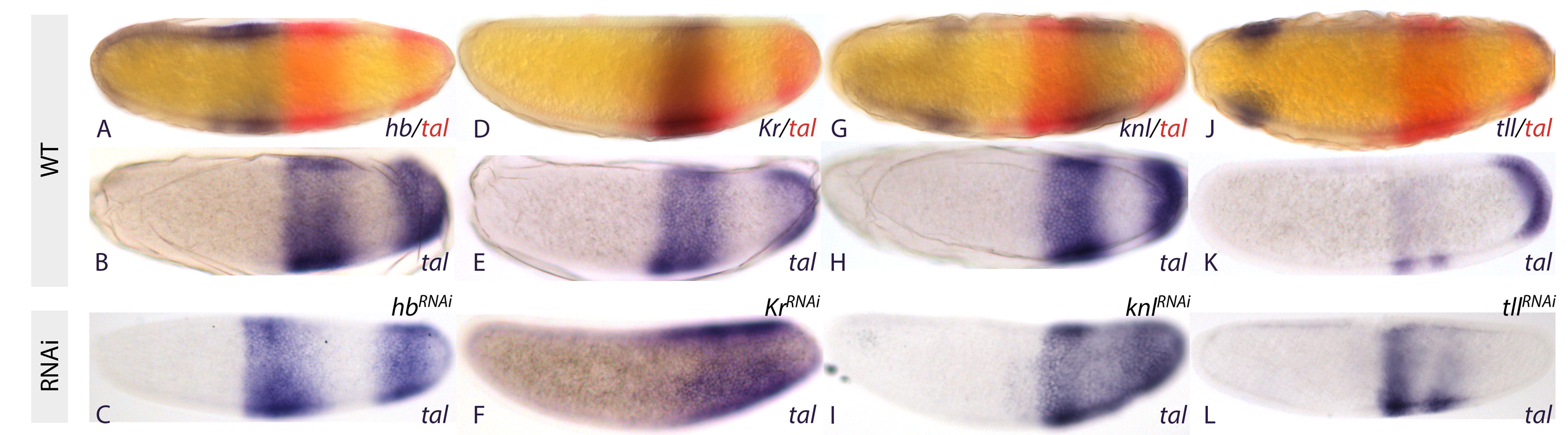
Effect of gap gene depletion by RNAi on *Ca-tal* expression. Colorimetric (enzymatic) *in situ* hybridisation against *Ca-tal* and gap genes in *C. albipunctata* blastoderm embryos. (A) Wild-type (WT) embryos stained against *Ca-tal* (red) and gap genes (blue) as indicated in the lower right corner of each panel. (B) Wild-type (WT) embryos stained against *Ca-tal* only (blue). (C) *Ca-tal* expression (blue) in embryos depleted for gap RNA as indicated in the top right corner of each panel. Embryos oriented laterally: anterior is to the left, dorsal to the top.

### Ca-tal *and* Ca-gt *do not exhibit gap gene phenotypes in* C. albipunctata

To further examine the function of *Ca-tal* and the gap genes in *C. albipunctata*, we obtained cuticle preparations of late-stage wild-type and RNAi embryos according to a previously published protocol (Figure 5) [22]. In cuticles of embryos treated with RNAi against Ca-hb (*n=22*), we observed a reduction in the number of thoracic segments in all specimens: seven embryos showed no, nine embryos one, and six embryos two remaining thoracic segments (Figure 5B). We only managed to obtain two cuticles of embryos treated with RNAi against *Ca-Kr.* Both of these exhibit general A–P polarity, but no thoracic or abdominal segments are discernible (Figure 5C). Similarly, severe defects were observed in the two cuticles we obtained from embryos treated with RNAi against *Ca-knl*: these embryos show two recognisable thoracic and one or two abdominal segments, albeit with severe dorsal defects, as well as an abnormal posterior terminal region (Figure 5D). Cuticles of embryos treated with RNAi against *Ca-tll* (*n=15*) show a much less penetrant phenotype. In four individuals, the telson is missing, and two show a severe reduction of the number of abdominal segments (Figure 5E, F). Only one specimen exhibited defects in the head and the thoracic region (not shown). We could detect no gap gene phenotypes or other obvious and consistent segmentation defects in embryos depleted for *Ca-tal* and *Ca-gt* (Figure 5G,H). However, in 2 out of 39 of the *gt* and 5 out of 55 of the *tal* depleted cuticles we observe small hemilateral abnormalities (Supplementary Figure 3 A, B, asterisks). We cannot rule out a weak effect of RNA depletion, but the cause of these abnormalities could also be mechanical or unspecific. We do not observe this type of effect in the other RNAi injected cuticles. The evidence from our cuticle preparations suggest that *hb, Kr, kni/knl*, and *tll* have conserved roles as gap genes in *C. albipunctata*, while *gt* and *tal* are expressed in a gap-like manner (*tal* also being regulated by other gap genes) but do not play a classical gap-like role in trunk segment determination in this species.

**Figure 5.**
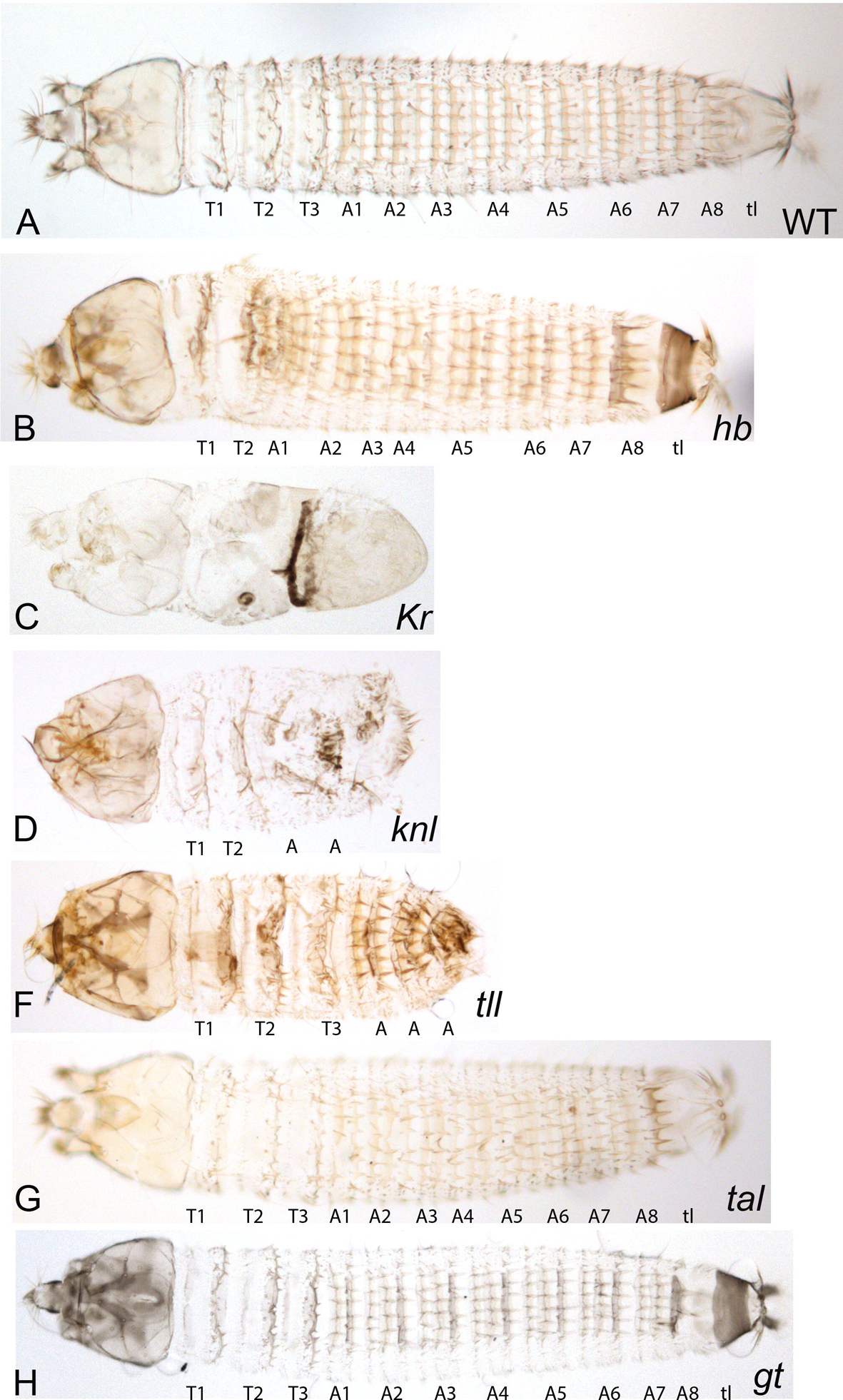
Cuticle preparations of late-stage wild-type and RNA-depleted embryos. (A) Wild type, (B) *Ca-hb* RNAi, (C) *Ca-Kr* RNAi, (D) *Ca-knl* RNAi, (E) *Ca-tll* RNAi, (F) *Ca-tal* RNAi and (G) *Ca-gt* RNAi. Labels indicate thoracic (T) and abdominal (A) segments, as well as the telson (tl). Wild-type cuticles include three thoracic segments, eight abdominal segments, and the telson (A). Segment numbers in depleted cuticles do not necessarily indicate exact segment identity, but reflect the number of segments present in each knock-down embryo. Cuticles shown in dorsal view, anterior is to the left. See Materials and Methods for details on preparation.

## Conclusions

We have characterised the homolog of *tal/pri/mlpt* in the nematoceran moth midge *C. albipunctata*. Similar to its homologs in other organisms [11,19], it produces a polycistronic primary transcript, which codes for several short peptides. We have shown that *Ca-tal* is expressed in a gap-gene-like manner in *C. albipunctata*, unlike in *D. melanogaster* where it initiates transcription in refined stripes during the late blastoderm stage [12]. We show that these early stages of expression are regulated by gap genes in *C. albipunctata.* Later expression patterns are more conserved between the two species. Despite its suggestive early embryonic expression pattern, *Ca-tal* cannot be classified as a segmentation gene. Our evidence reveals that *Ca-tal* is not regulating other segmentation genes, and does not cause a gap-like or any other segmentation phenotype upon knock-down by RNAi.

The gap-like expression pattern of *Ca-tal* shows striking similarities to its homolog, the gap gene *mlpt* in the flour beetle *T. castaneum.* However, even this similarity may be superficial, as there are significant differences between the regulation of both homologs. The anterior domain of *Ca-tal* is repressed by Hb, while the posterior terminal domain is not affected in *hb* RNAi knock-downs (summarized in Figure 6). In *T. castaneum*, the opposite is true: while the anterior domain of *mlpt* is not affected, the posterior domain forms late in if *hb* is depleted [19]. Furthermore, *Ca-tal* is repressed by Knl (Figure 6), while Kni does not affect *mlpt* expression in *T. castaneum* [19]. In contrast, *mlpt* is activated by Gt [19], while *Ca-tal* and *gt* show no genetic interaction. The only similarity between the two species is the role of *Kr* in *tal/mlpt* regulation: ectopic expression is seen upon *Kr* knock-down in the posterior of blastoderm embryos in *C. albipunctata* and *T. castaneum.* Based on the available evidence, it remains unclear whether the early gap-like expression pattern of *tal/mlpt* is an ancestral feature of segmentation patterning, or whether it has evolved convergently in beetles and nematoceran dipterans. Functional data from other basally branching dipteran lineages or suitable outgroups will be required to resolve this outstanding question.

**Figure 6.**
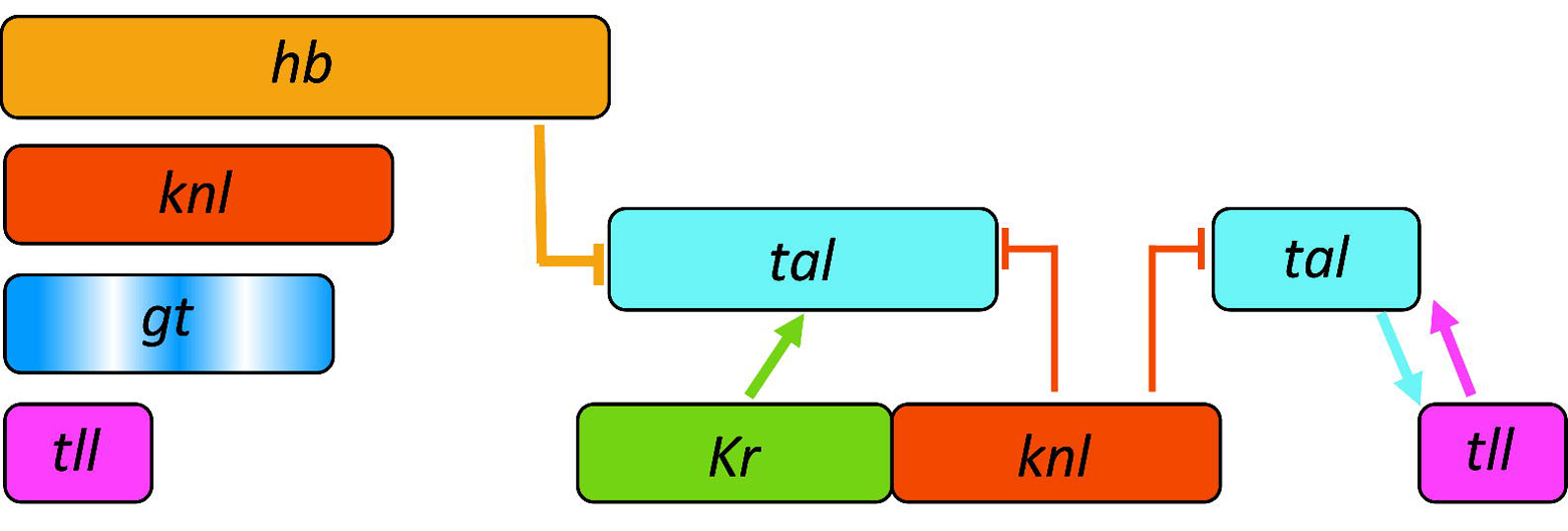
Schematic summary of *Ca-tal* expression and regulation. Coloured boxes indicate the relative position of expression domains along the A–P axis (anterior is to the left, posterior to the right). Arrows indicate activation, T-bars indicate repression, as suggested by our RNAi knock-down experiments.

Supplementary Figure 1. Effect of *Ca-tal* depletion by RNAi on gap gene expression. Colorimetric (enzymatic) *in situ* hybridisation in wild type (A–F) and *Ca-tal* depleted embryos (G–L) against *Ca-hb* (A, G), *Ca-gt* (B, H), *Ca-Kr* (C, I), *Ca-knl* (D, J), and *Ca-tll* (E, F, K, L). (L, *tll\**) shows an example for the 4 out of 17 embryos where posterior expression of *Ca-tll* is repressed (marked by an asterisk) in embryos depleted for *Ca-tal* by RNAi; in most embryos, posterior *Ca-tll* expression is not affected (K). Embryos oriented laterally: anterior is to the left, dorsal to the top.

Supplementary Figure 2. Lack of effect of *Ca-Kr* depletion on *Ca-knl* expression. Colorimetric (enzymatic) *in situ* hybridisation against *Ca-knl* (blue) in wild-type (WT, A), and *Ca-Kr* depleted embryos (B). Wild-type embryo in (A) is co-stained against *Ca-tal* (red). There are no detectable differences in *Ca-knl* expression between wild-type and *Ca-Kr* knock-down. Embryos oriented laterally: anterior is to the left, dorsal to the top.

Supplementary Figure 3. Cuticle preparations of late-stage RNA-depleted embryos for (A) *Ca-gt* RNAi, and (B) *Ca-tal* RNAi. Both example embryos show hemilateral cuticle abnormalities (indicated by asterisks). Frequencies of these knock-down phenotypes are very low (see text for details). Cuticles shown in dorsal view, anterior is to the left. See Materials and Methods for details on preparation.

## Materials and Methods

### Gene identification, cloning, and synthesis of RNA constructs

We searched the early embryonic transcriptome of *C. albipunctata* ([20]; http://diptex.crg.es) using peptide sequences from *D. melanogaster tarsal-less* retrieved from GenBank. A 2277 nt clone was obtained (Diptex clone CAL_comp2583_c0_seq1). This sequence was confirmed via PCR on *C. albipunctata* cDNA, and is deposited in GenBank under accession number MG783326. A 1.2kb fragment of *Ca-tal*, containing all small ORFs, was used to obtain riboprobes for whole-mount *in situ* hybridisation, as well as template for double-stranded RNA. For the gap genes, clones were the same as used in [9] (fragment size in parentheses): *Ca-hb*, AJ131041.1 (1800nt); *Ca-Kr*, GU137323.1 (1200nt); *Ca-knl*, GU137321.1 (800nt); *Ca-gt*, GU137318.1 (1100nt); *Ca-tll*, GU137320.1 (1400nt). Fragments were cloned into the PCRII-TOPO vector (Invitrogen), and used to make DIG or FITC-labelled riboprobes for whole-mount *in situ* hybridisation, as well as double-stranded RNA. RNAi constructs were synthesised as described in [22].

### Embryo collection and fixation

Wild-type and RNAi-treated embryos of *C. albipunctata* were collected at blastoderm and post-gastrulation stages as described previously in [9]. Embryos were heat-fixed using a protocol adapted from [24]. In brief, embryos were dechorionated at the desired developmental stage by immersing them in 25% bleach for 45s. Embryos where then dipped in boiling fixing solution (0.7% NaCl; 0.05% Tween20) for 20s. Heat fixation was stopped by adding room-temperature (RT) water to the solution. Embryos were subsequently post-fixed in formaldehyde (5%) and PBS/methanol. Devitellinisation is achieved by vigorous shaking for 20s in 50% heptane-methanol. Embryos were preserved in methanol. For RNAi-treatment, embryos were dechorionated manually using tungsten needles and fixed as described for wild-type.

### Whole-mount *in situ* hybridisation

Whole-mount *in situ* hybridisation was performed as described for *C. albipunctata* in [22] and references therein. In brief, embryos were permeabilised after rehydration using Proteinase K (8 mg/ml PBT) for 7 min at RT, followed by post-fixation in 5% formaldehyde/PBT for 25 min. Overnight hybridisation at 56°C was carried out with a labelled probe at a concentration of 0.5–1 ng/μl. For detection, antibodies (anti-digoxigenin or fluorescein conjugated with alkaline phosphatase (Sigma)) were used at 1:2000 for 2h. Staining was achieved using NBT/BCIP (blue) or fast red. Embryos were counterstained with DAPI, and mounted in 70% glycerol.

### RNA interference (RNAi)

RNAi treatment was carried out based on protocols established in other dipteran species [4,5,25] as described previously in [22]. In brief, embryos were allowed to develop for 2h before immersing them in 25% bleach for 10s to weaken the chorion, then rinsed under tap water for 1 min. Embryos where aligned on a microscope slide against a glass capillary (Hilgenberg 1421602, 65mm x OD 0.25mm) and covered with a 3:1 mixture of 10:27 halocarbon oil. Alumniosilicate (rather than borosilicate) capillaries (pulled in Sutter P-97 Flaming/Brown Micropipette Puller) were used for the injections, maintaining a constant flow of liquid to avoid blocking of the needle. After injection embryos were allowed to develop for 7h before being fixed and collected as previously described [24]. Buffer-only injections were used as a negative control, along with *in situ* hybridisation staining for depleted transcripts. Cuticle preparations were performed as described in [22]. Only cuticles injected with double-stranded RNA presented abnormal phenotypes. Double-stranded RNA was injected at the following concentration: *Ca-hb*, 5.1 μM; *Ca-gt*: 5.9 and 3.8 μM; *Ca-Kr*: 5.4 μM; *Ca-knl*: 5.1 μM, *Ca-tal*: 7.9 and 4.2 μM; Ca-*tll*: 3.4 μM.

## Ethics

The animals used in this study do not require any ethical approval.

## Competing interests

We have no competing interests.

## Authors’ contributions

EJ-G designed the study, performed experiments, and wrote the paper. KRW performed RNAi experiments, and contributed to writing the paper. JJ conceived and designed the study, financed the experiments, and contributed to writing the paper. All authors gave final approval for publication.

## Acknowledgements

The authors would like to thank Damjan Cicin-Sain for help and support with the FlyGUI/FlyAGE image-processing pipeline, Nuria Bosch for help with the fly culture, and Isma de Mingo for informatics technical support. We thank Juan Pablo Couso and Urs Schmidt-Ott for inspiring and encouraging critical discussions at early stages of this project.

## Funding

This work was funded by the MEC-EMBL agreement for the EMBL/CRG Research Unit in Systems Biology, SGR Grant 406 from the Catalan funding agency AGAUR, and by grants BFU2009-10184 and BFU2012-33775 from the Spanish Ministerio de Economia y Competitividad (MINECO). The Centre for Genomic Regulation (CRG) acknowledges support from MINECO, “Centro de Excelencia Severo Ochoa 2013-2017”, SEV-2012-0208.

